# Interferon-Gamma Signaling Promotes Melanoma Progression and Metastasis

**DOI:** 10.1101/2021.10.14.464463

**Authors:** Bo Zhou, Jayati Basu, Hasan R. Kazmi, Xuan Mo, Sarah Preston-Alp, Kathy Q. Cai, Dietmar Kappes, M. Raza Zaidi

**Affiliations:** Fels Cancer Institute for Personalized Medicine, Lewis Katz School of Medicine, Temple University, Philadelphia, PA, USA; Fox Chase Cancer Center, Philadelphia, PA, USA; Janssen Pharmaceutical Companies of Johnson & Johnson, Spring House, PA, USA

## Abstract

Interferon-gamma (IFNG) has long been regarded as the flag-bearer for the anti-cancer immunosurveillance mechanisms. However, relatively recent studies have suggested a dual role of IFNG, albeit there is no direct experimental evidence for its potential pro-tumor functions. Here we provide *in vivo* evidence that treatment of mouse melanoma cell lines with physiological levels of Ifng enhances their tumorigenicity and metastasis in lung colonization allograft assays performed in immunocompetent syngeneic host mice, but not in immunocompromised host mice. We also show that this enhancement is dependent on downstream signaling via Stat1 but not Stat3, providing evidence of an oncogenic function of Stat1 in melanoma. The experimental results suggest that melanoma cell-specific Ifng signaling modulates the tumor microenvironment and its pro-tumorigenic effects are dependent on the γδ T cells, as Ifng-enhanced tumorigenesis was inhibited in the TCR-δ knockout mice. Overall, these results show that Ifng signaling may have tumor-promoting effects in melanoma by modulating the immune cell composition of the tumor microenvironment.

## INTRODUCTION

Cutaneous malignant melanoma is a complex, highly aggressive, and frequently chemoresistant cancer that continues to exhibit a positive rate of increase in the developed world (Gandini et al., 2005; Nikolaou and Stratigos, 2014; Tran et al., 2008). Numerous epidemiological studies have identified the solar ultraviolet radiation (UV/UVR) to be the major etiological risk factor for melanoma (Garibyan and Fisher, 2010; Garland et al., 2003; Moan et al., 2008), with the highest risk associated with intermittent burning doses, especially during childhood (Austin et al., 2013; Bennett, 2008; Maddodi and Setaluri, 2008; Slade and Austin, 2014; Whiteman et al., 2001). However, the underlying molecular mechanisms by which UVR (UVB and UVA wavebands) initiate melanomagenesis remain poorly understood. Although numerous studies have amassed strong evidence that UVB-induced signature DNA mutations play key roles in melanomagenesis, there is compelling evidence that non-mutational mechanisms, such as UVR-induced inflammation and immunosuppression, also contribute substantially to melanomagenesis (Hocker and Tsao, 2007; Matsumura and Ananthaswamy, 2002; Norval et al., 2008), highlighting the importance of mechanisms other than direct DNA damage in UVR-induced initiation and/or progression of melanoma.

To uncover the molecular mechanisms underlying UVR-induced melanomagenesis, we previously investigated the genomic response of melanocytes to UVB and UVA radiation (Zaidi et al., 2011). We showed that answers to many of the questions regarding UVR-induced melanomagenesis lie not only in how UVR damages melanocyte DNA but also in how altered gene expression in the exposed melanocytes drives their interactions with the elements of the microenvironment to remodel damaged skin and allows UVR-damaged (mutated) melanocytes escape immunosurveillance-based destruction. We showed that UVB can directly upregulate melanocytic expression of ligands to the chemokine receptor CCR2, which recruits macrophages into the neonatal skin microenvironment. A subset of these macrophages produces interferon-gamma (IFNG) into the tumor microenvironment, which we postulated to be paradoxically pro-melanomagenic (Zaidi et al., 2011).

IFNG is known to play a central role in cancer immunosurveillance and immunoediting (Ikeda et al., 2002; Schreiber et al., 2011). IFNG is also associated with anti-proliferative and anti-angiogenic, as well as anti-tumor immune responses against a variety of different cancers, including melanoma (Brown et al., 1987; Dummer et al., 2004; Ikeda et al., 2002; Tamura et al., 1987; Wall et al., 2003). For example, it has been reported that IFNG had significant growth inhibitory activity on four different human melanoma cell lines; albeit at concentrations that were 1000- to 10,000-fold higher than physiologic levels (Kortylewski et al., 2004). However, a potentially pro-tumorigenic role of IFNG has been postulated (Zaidi, 2019; Zaidi and Merlino, 2011). There is some indirect evidence that indicates that IFNG can have contrasting roles in tumorigenesis, i.e. it can exhibit both cytostatic/cytotoxic and anti-tumorigenic immunosurveillance functions as well as pro-tumorigenic immune-evasive effects in the tumor microenvironment in a context-dependent manner. Here we report in vivo evidence that activation of IFNG signaling directly in melanoma cells enhances their tumorigenicity and metastasis, which is dependent on the melanoma cell-driven modulation of the tumor immune microenvironment.

## RESULTS

### Ifng treatment of mouse melanoma cells enhances lung colonization and metastasis

To understand the effects of IFNG signaling on melanoma cells, we studied mouse melanoma cell growth *in vitro* with and without treatment with physiological levels (10 ng/ml) of recombinant Ifng. The mouse melanoma cell lines tested showed differential response to Ifng. The B16 and its derivative cell line B16N showed significant reduction in proliferation (Figures S1A and S1B). Surprisingly, however, the proliferation of B2905, F5061, and YUMM1.1 cell lines was not affected by Ifng treatment, as no significant differences were seen in proliferation (Figures S1C, S1D, and S1E). Similarly, while the soft agar colony formation of B16 and B16N cells was significantly inhibited by Ifng treatment (10 ng/ml) (Figures S2A and S2B), B2905 and F5061 cell lines did not show any difference in colony formation after Ifng treatment (Figures S2C and S2D).

To assess the role of Ifng on melanoma tumorigenesis in mouse allograft model systems, we pretreated 5 mouse melanoma cell lines (B16, B16N, B2905, F5061, and YUMM1.1) in culture for 48 h with 10 ng/ml Ifng and implanted them into immune-competent syngeneic mice (C57BL/6 or FVB/N) via subcutaneous or tail-vein injections (Figure 1A). While no differences were detected in subcutaneous tumorigenesis between the mock-treated control and Ifng-treated groups (Figure S3), the tail-vein inoculation models showed a striking increase in the number of tumor nodules on the lung surfaces and/or by histopathological analyses of the lung tissues in the mice harboring Ifng-treated cells of all 5 melanoma lines (Figures 1B, 1C, 1D, 1E, 2A, and S4). The pigmented lung nodules of the B16, B16N, and B2905 cells could be observed both visually and histologically (Figures 1B and 1C), whereas those of F5061 and YUMM1.1 cells were unpigmented and were only analyzable by histological analyses (Figures 1D and 1E). To assess the effect of Ifng treatment on the metastatic potential of the B16N cells (a new metastatic clone of B16), we injected Ifng-treated and control B16N cells with ectopic expression of luciferase via tail vein in syngeneic C57BL/6 mice and monitored tumor growth by bioluminescence imaging. We observed a significant increase in tumor growth in mice harboring Ifng-treated cells as compared to the controls, as measured by bioluminescence (Figures 2B and 2C). Posthumous analyses showed a statistically significant increase in extrapulmonary metastases in the ovary, bones, parametrium, and kidney tissues of the mice inoculated with Ifng-treated cells as compared with the mock-treated controls (Figure 2D).

**Figure 1.**
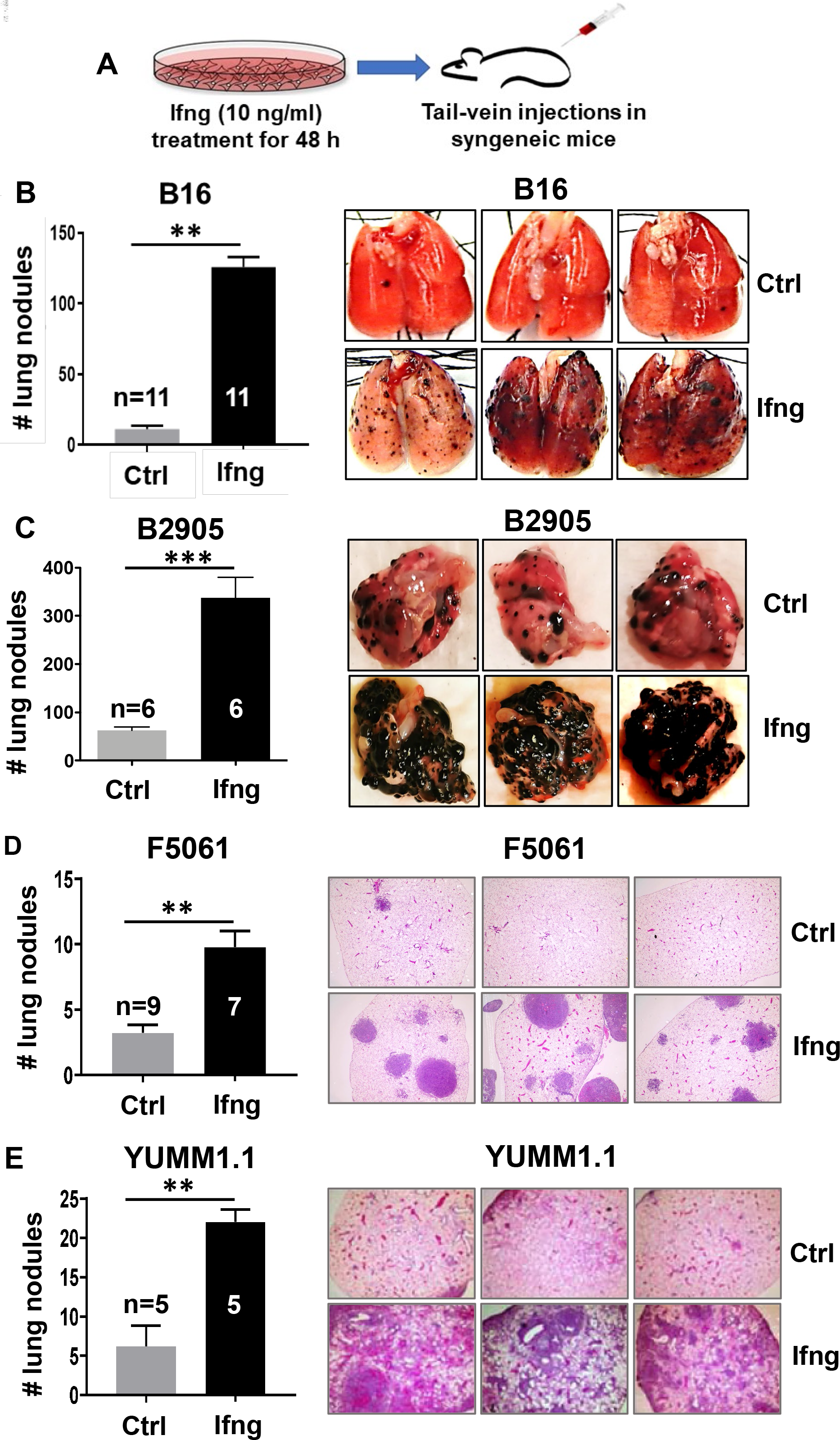
Induction of Ifng signaling enhances melanoma tumorigenesis. (A) An illustration of the experimental plan is shown. Mouse melanoma cell lines were treated with 10 ng/ml recombinant mouse Ifng (or mock treated, Ctrl) for 48 h, followed by inoculation in syngeneic mice via tail vein injections. (B-E) Quantification of tumor nodules in lungs, either by counting surface nodules on gross lungs (for pigmented tumors) or in H&E-stained sections from paraffin-embedded lung tissues (for unpigmented tumors). Significantly increased tumor counts were observed in all mice inoculated with Ifng-treated cell lines. Lungs recovered 20-28 days after injection from mice bearing (B) B16 and (C) B2905 cells were paraformaldehyde-fixed, quantified, and photographed using a dissecting microscope. Pulmonary tumors appeared as black pigmented nodules on the surface of the lungs. For the unpigmented cell lines, (D) F5061 and (E) YUMM1.1, representative photomicrographs of 4x microscopic fields of view in H&E-stained sections are shown. 3 slides per mouse, 3 fields per slide, for a total of ≥45 fields were quantified per group. Data are presented as mean±SEM. n = number of mice used for each group. *P< 0.05, **P<0.01, ***P<0.001), Student’s t test.

**Figure 2.**
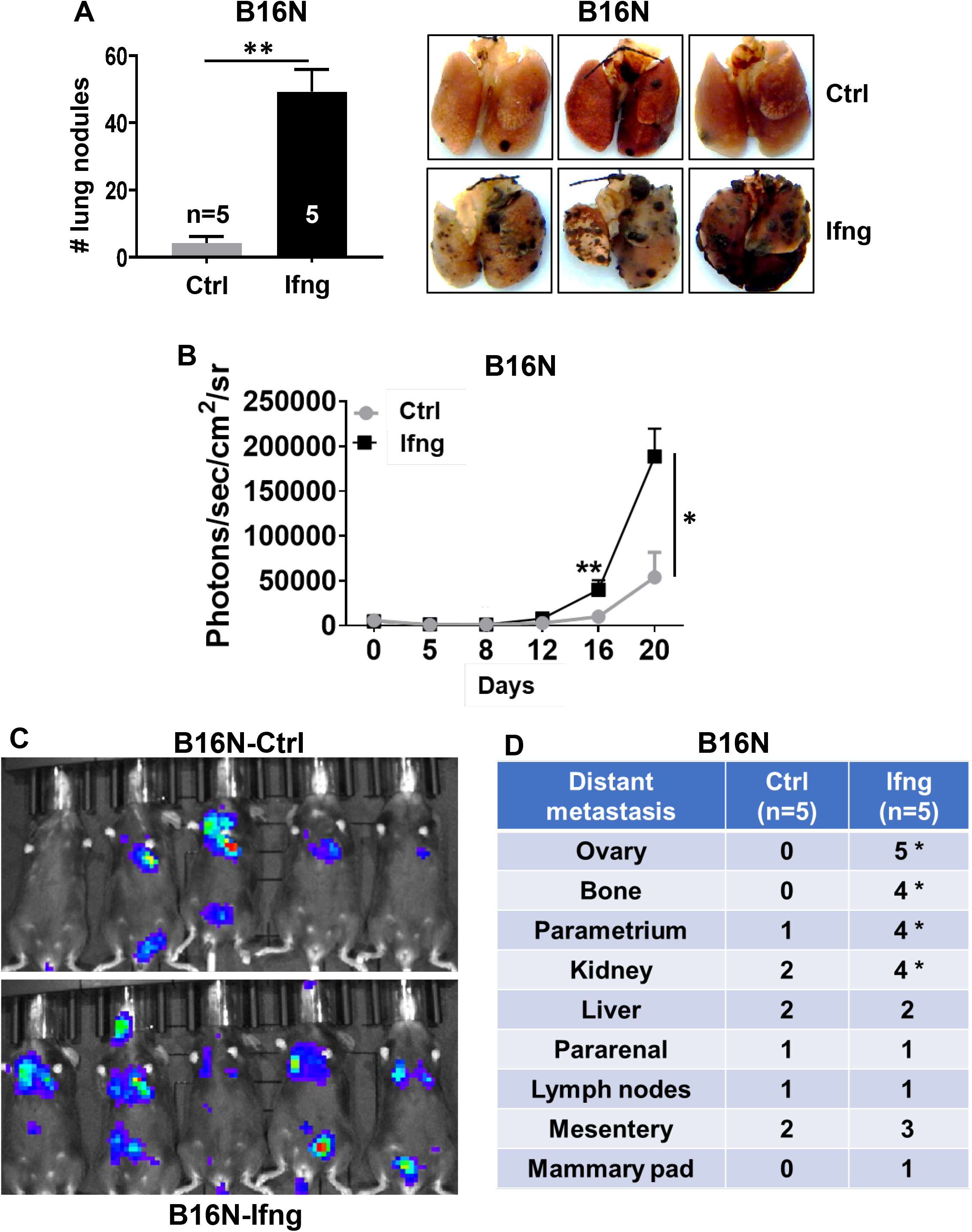
Ifng signaling enhances metastasis. Bioluminescent B16N metastatic melanoma cell line was treated (or mock treated, Ctrl) with Ifng (10 ng/ml) for 48 h, followed by tail vein injection into syngeneic C57BL/6 mice. Tumor growth was monitored by live animal bioluminescence imaging at days 5, 8, 12, 16 and 20. Lung surface nodules were counted at terminal autopsy on day 20. (A) Quantification of lung surface tumor nodules. n = 5 mice each group. Representative lungs are shown. Data are presented as mean±SEM. **P<0.01, Student’s t test. (B) Total tumor burden as measured by bioluminescence (photon flux) imaging in live mice. Metastasis by Ifng-treated B16N cells was significantly increased compared to the control cells. *P<0.05, **P<0.01, Student’s t test. (C) Bioluminescence imaging on day 20 after cell inoculation. (D) The frequency and distribution pattern of extrapulmonary metastases was determined in the indicated tissue samples collected at 20 d after tail vein injection by a detailed histopathological analysis. The frequency of extrapulmonary metastases was significantly greater (*p<0.05, chi-square test) in the mice inoculated with Ifng-treated B16N cells.

### Stat1 but not Stat3 is the downstream effector of the pro-tumorigenic effects of Ifng signaling

The activation of the canonical Ifng signaling pathway leads to the phosphorylation of Jak1 and Jak2, followed by phosphorylation and homodimerization of the transcription factor Stat1, which translocates to the nucleus to activate its target genes. However, in some contexts, Ifng can also activate Stat3 (Qing and Stark, 2004). We observed phosphorylation of both Stat1 and Stat3 upon Ifng treatment in mouse melanoma cell lines. To determine whether the Ifng-mediated tumorigenic effect on melanoma cells was routed through Stat1 or Stat3, we generated knockout (KO) clones for both Stat1 and Stat3 in the B16N cell line utilizing the CRISPR/Cas9 methodology. The knockout clones were verified by western blotting to confirm the absence of the respective proteins (Figure S5). The B16N-Stat1-KO and B16N-Stat3-KO cells were treated with 10 ng/ml Ifng or mock-treated as above and their proliferation and colony formation were assessed. The results showed that while the parental WT B16N cells and their Stat3-KO counterparts were significantly inhibited in proliferation by Ifng treatment, this effect was nullified in the Stat1-KO cells (Figure S6A). In the colony formation assay, Stat1-KO cells did not respond to Ifng treatment; however, colony formation was suppressed in Stat3-KO cells (Figures S6B and S6C).

To test the in vivo effects of Stat1-KO and Stat3-KO, cells were inoculated via tail vein in syngeneic mice. While the Ifng-treated B16N-Stat3-KO cells showed a statistically significant increase in lung tumorigenesis as compared to the controls (Figure 3A), which was similar to the parental B16N cells, the tumorigenicity of the B16N-Stat1-KO cells was drastically and statistically significantly inhibited and there was no difference in lung colonization between Ifng-treated and control B16N-Stat1-KO cells (Figure 3B). These results suggested an obligate requirement of Stat1 but not Stat3 as the downstream mediator of the tumorigenic effects of Ifng signaling on melanoma cells.

**Figure 3.**
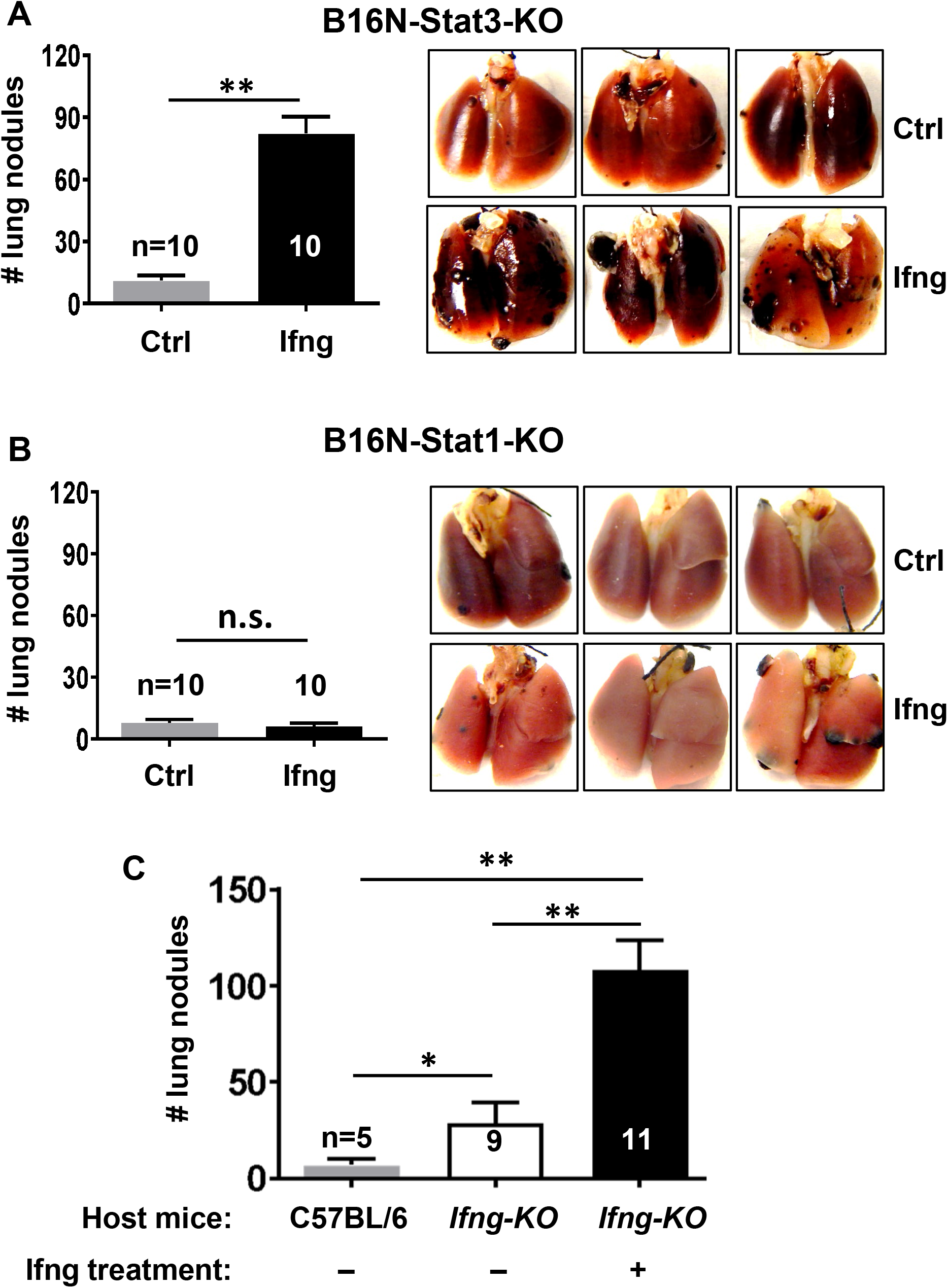
Ifng-mediated enhancement of lung tumorigenesis is through Stat1 but not Stat3. (A) Stat3-knockout (Stat3-KO) and (B) Stat1-KO B16N cells were treated with 10 ng/ml Ifng (or mock treated, Ctrl) and injected by tail vein into C57BL/6 mice. Lung surface tumor nodules were quantified at 20 d. n = 10 each group. n.s. = not significant; **P<0.01, Student’s t test. (C) Ifng signaling in melanoma cells enhances tumorigenesis independently of the host systemic Ifng signaling, which is anti-tumorigenic. Untreated B16N cells were tail vein inoculated in either wild type C57BL/6 (n = 5) or C57BL/6-*Ifng-knockout* host mice (n = 9). The untreated cells exhibited significantly enhanced tumorigenesis in the *Ifng-KO* host mice. Ifng-treated B16N cells showed further enhancement of tumorigenesis in Ifng-KO host mice (n = 11). Data are presented as mean±SEM of lung surface tumors. *P<0.05, **P<0.01, Student’s t test.

To determine the comparative contribution of the systemic Ifng signaling of the host mice and the intracellular Ifng signaling in melanoma cells to the enhancement of tumorigenicity, we injected untreated parental B16N cells in wildtype C57BL/6 and *Ifng-knockout* (C57BL/6) host mice. There was a statistically significant increase in tumorigenicity of the cells in the *Ifng-KO* host mice, confirming the essential role of Ifng in the systemic anti-tumor immunosurveillance mechanisms. Remarkably, a further additive increase in the cells’ tumorigenicity was seen when the cells were treated with Ifng followed by injection in *Ifng-KO* host mice (Figure 3C). These results suggest dual and opposite functions of Ifng signaling wherein it plays an anti-tumor role in the context of the systemic immunosurveillance but has a pro-tumor effect on the melanoma cells.

### Ifng-enhanced melanoma tumorigenicity is dependent on the immune system

To test whether the enhancement of tumorigenicity and metastatic capabilities of melanoma cells by Ifng is dependent on the presence of the immune system, we implanted B16N and B2905 cells, with or without treatment with Ifng, in the immunocompromised NOG mice via tail vein (Figure 4A and 4B). There were no statistical differences in lung colonization between the treated and control groups, indicating that the Ifng-mediated enhancement of tumorigenesis required a functional immune system. Similar insignificant results were obtained when a human melanoma cell line A2058, with or without treatment with IFNG, was inoculated in the NOG mice via tail vein (Figure 4C and 4D). To test whether the absence of the T cell compartment was responsible for the lack of Ifng-mediated enhancement of tumorigenesis in the immunocompromised context, we inoculated Ifng-treated and control B16N cells in the athymic Nude (*Foxn1*^*nu*^) mice via tail vein. Interestingly, we observed a reduced number of lung tumor nodules but robust extrapulmonary metastatic spread, both of which were not statistically different between the Ifng-treated and control groups (Figure 4E). These results implicated the modulation of the T cell-mediated immunity as the central player in the Ifng-mediated enhancement of melanoma tumorigenesis.

**Figure 4.**
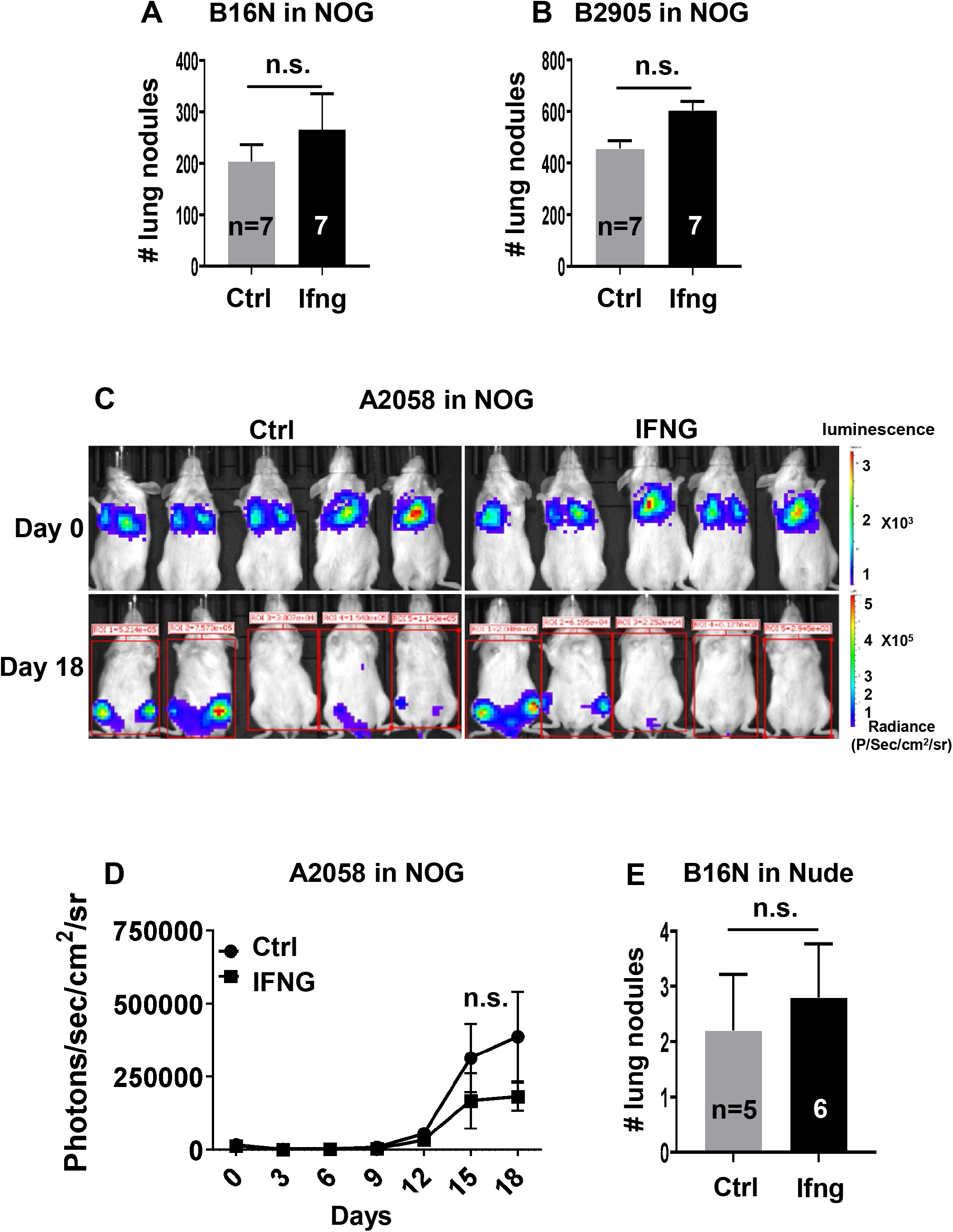
Host immune response is necessary for the Ifng-mediated enhancement of melanoma lung colonization. Mouse melanoma cells (A) B16N and (B) B2905 were treated with 10 ng/ml of Ifng and inoculated into the immunodeficient NOG mice (n = 7 for each group) via tail vein. Quantification of lung surface tumor nodules is shown as mean±SEM. n.s. not significant; Student’s t test. (C) A2058 human melanoma cells were treated (or mock treated, Ctrl) with human recombinant IFNG and injected into NOG mice through the tail vein and monitored by bioluminescence imaging (BLI). BLI images at day 0 immediately after tail vein injection and on 18 d are shown. (D) Total tumor burden (photon flux) as measured by BLI. n.s. not significant, One-Way ANOVA with post-hoc analysis. (E) Ifng-treated (n = 6) and Ctrl (n = 5) B16N cells were tail vein inoculated in the immunocompromised Nude mice. Lung surface tumor nodules were quantified at 18 d post inoculation. n.s. not significant; Student’s t test.

### Ifng-treated melanoma cells modulate the tumor immune microenvironment

Since, we found no difference in lung tumorigenesis of Ifng-treated versus untreated melanoma cells when inoculated in T cell-deficient hosts, we hypothesized that Ifng-treated melanoma cells modulate their tumor microenvironment where T cells play a crucial role. To compare the tumor microenvironment (TME) between untreated versus Ifng-treated melanoma lung tumor nodules, we microdissected the tumor nodules from the lungs and performed immune phenotyping by multi-color flow cytometry. Interestingly, the tumor nodules formed by the Ifng-treated melanoma cells and the tumors from the control melanoma cells did not show a statistically significant difference in the tumor-infiltrating CD45+ cells in the TME (Figure 5A). We found no significant differences in the frequencies of tumor-infiltrating lymphocytes (TILs), CD4, CD8, and T_reg_ cells between the two groups (Figures 5B, 5D, and 5E. However, further analyses revealed a statistically significant increase in CD4+CD8+ dual-positive (DP) T cells (Figure 5D and 5I). Surprisingly, the frequency of tumor-associated neutrophils (TAN) was increased >2-fold in the Ifng-treated melanoma TME (Figures 5C and 5I). In the γδ T cell compartment, although the frequency of the infiltrating γδ T cells was similar, we observed a significant increase in the CD27-CCR6-γδ T cells with a concomitant decrease in the CD27+ γδ T cells in the Ifng-treated melanoma TME as compared to the control tumors (Figures 5B, 5F, 5I, and 5K). The CD27+ γδ T cell are well known to produce Ifng, whereas the CCR6+ γδ T cells produce IL-17 and are known to perform a pro-tumorigenic role (Ribot et al., 2009). Therefore, we tested the cytokine production from γδ T cells, CD4, and CD8 T cells. As expected, γδ T cells from the Ifng-treated melanoma TME failed to produce Ifng (Figures 5H and 5K); whereas they readily produced both TGFβ and IL-17 upon in vitro stimulation of PMA and Ionomycin (Figures 5H and 5L). We also observed a trend of harboring a reduced frequency of Ifng-producing CD4+ cells in the TME in the Ifng-treated tumors (Figure 5G). Altogether, our analyses suggested that Ifng-treated melanoma cells created a TME that was enriched in neutrophils and IL-17- and TGFβ-producing γδ T cells.

**Figure 5.**
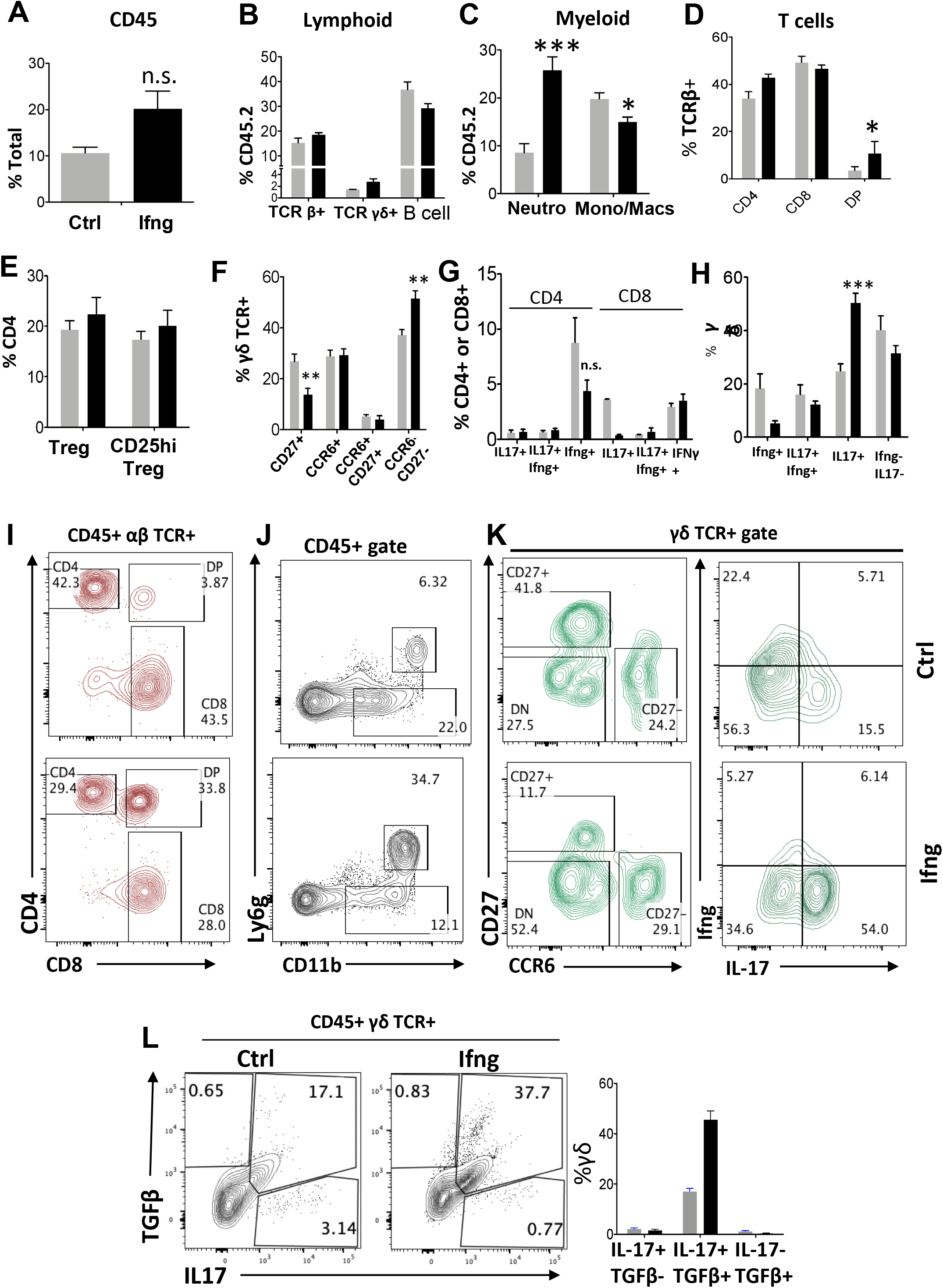
Analyses of immune cell infiltration in the tumor microenvironment. Lung tumors were isolated by microdissection from Ifng-treated and untreated B2905 tumor bearing mice at 23 d post inoculation. Tumor infiltrating immune cell profiles were generated by flow cytometric analysis. Relative cell frequencies within the indicated gated immune cell subset are shown for total lymphocytes (A), B cells (B), αβ T cells (B,D,I), γδ T cells (B,F,H,K) and myeloid cells (C, J). Cytokine production of infiltrating T cells in response to PMA and Ionomycin stimulation was measured by intracellular staining (G, H, K, L). Data displayed in panels A-H represent mean ± SEM. Significance of differences was determined by One-Way ANOVA with post-hoc Bonferroni test, as marked by: *P< 0.05; **P< 0.01; ***P< 0.001; ****P< 0.0001.

### Ifng-enhanced metastasis of melanoma cells is dependent on gamma-delta T cells

To further study whether the γδ T cells play a pivotal role in shaping a strong pro-tumorigenic microenvironment as opposed to the αβ T cells, we inoculated via tail-vein Ifng-treated and control B2905 melanoma cells in TCRδ−/− and TCRβ−/− mice. In contrast to WT and TCRβ−/− host mice, we observed a statistically significant reduction in lung colonization and metastasis of Ifng-treated cells in the TCRδ−/− mice (Figures 6A and 6B). In addition, the TCRδ−/− host mice exhibited a statistically significant reduction in the metastasis of the untreated control cells. A flow cytometry analysis of the TME revealed that lack of γδ T cells caused a substantial reduction in neutrophil infiltration in the TME of the tumors made by both Ifng-treated and untreated control cells (Figures 6C and 6D). A deficiency of the γδ T cells enhanced the infiltration of B cells in the TME (Figures S7A, S7B, S7C, S7D, and S7E), though we did not find any effect on the total number of T cells as CD4+, and CD8+ T cells in both groups (Figure S7B). Interestingly, we observed a marked decrease in the CD25-hi FoxP3+ T_regs_ in the Ifng-treated TME in the TCRδ−/− mice, as compared to WT, though the frequency of total Treg in TME did not change significantly (Figures 6F and 6G). Further analyses of the cytokine profile revealed that Ifng-producing CD4+ T cells were substantially increased in the Ifng-treated TME in the TCRγδ−/− host mice, as compared to WT (Figures 6H and 6I). We also observed a lack of γδ T cells-enhanced Ifng production by CD8+ TILs (Figure 6J). Collectively, our data suggest that Ifng signaling in melanoma cells remarkably enhances lung colonization and tumorigenesis via driving the recruitment of IL-17 and TGFβ-producing γδ T cells to enhance neutrophil recruitment to create a pro-tumorigenic microenvironmental niche.

**Figure 6.**
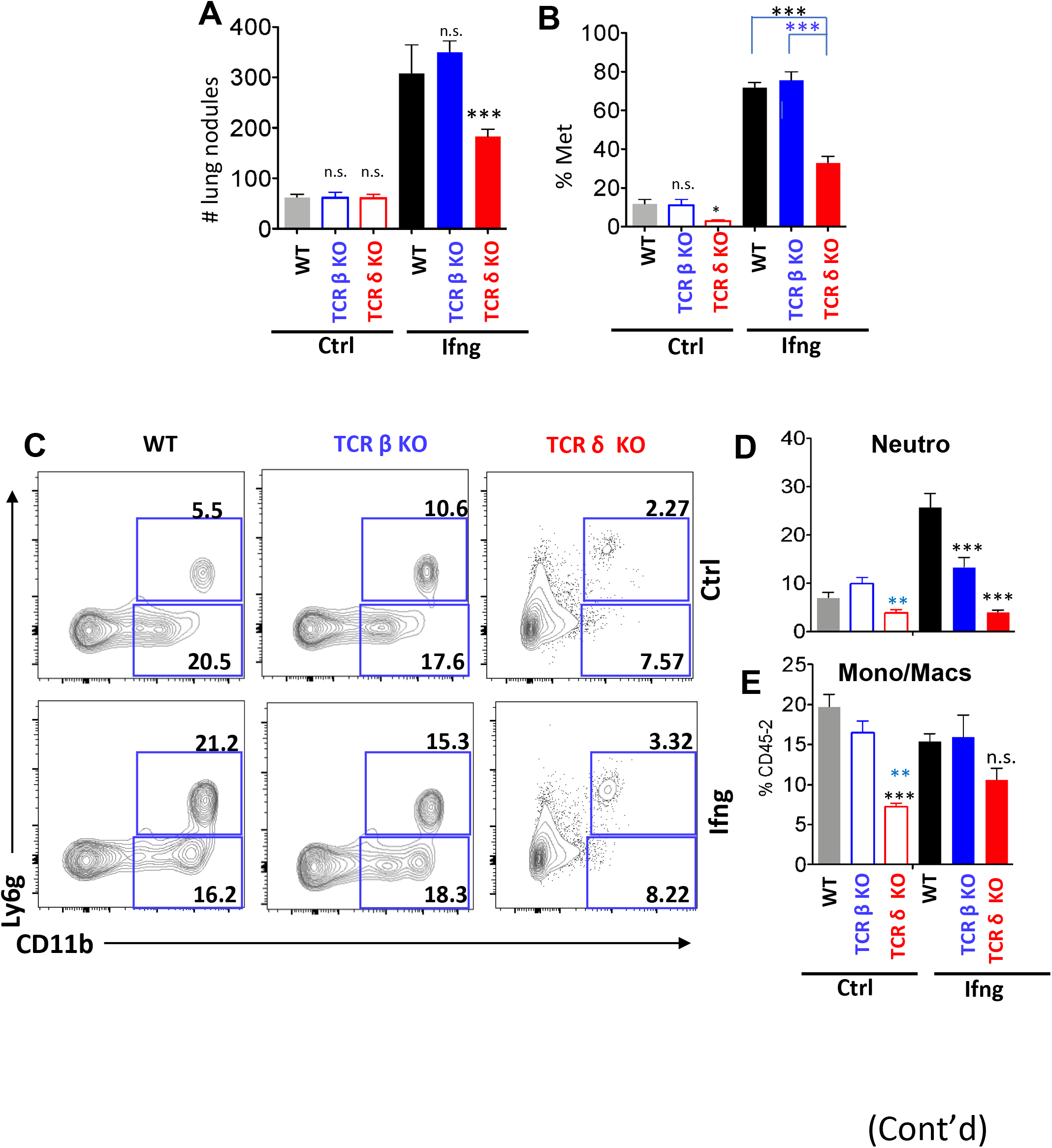
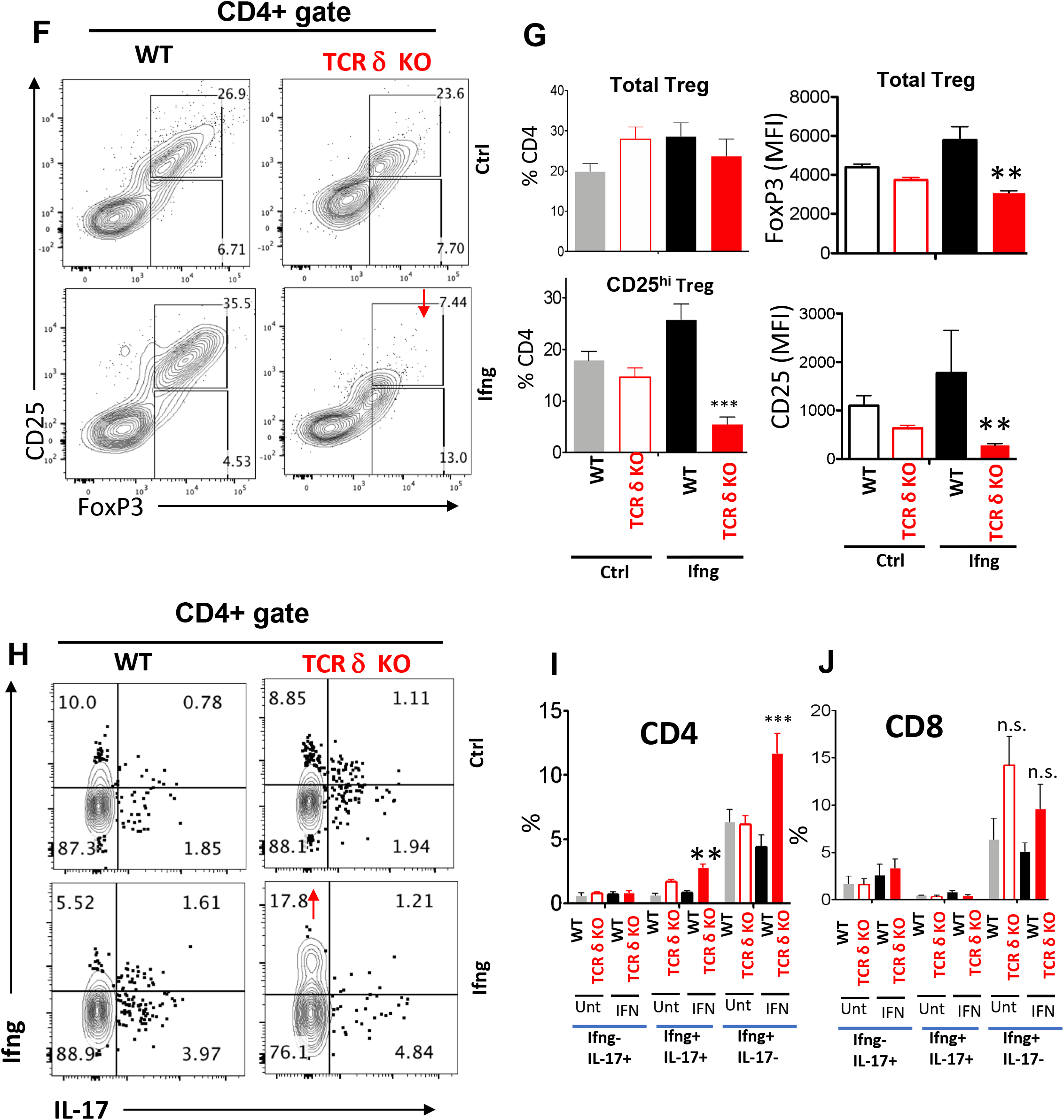
Requirement for gamma-delta T cells in Ifng-enhanced melanoma lung tumorigenesis. Ifng-treated and untreated B2905 melanoma cells were inoculated into syngeneic wild type, TCRβ−/− and TCRγ−/− mice by tail vein inoculation. (A) Mice were euthanized on 23 d and lung surface tumors were counted. (B) The proportion of lungs affected by metastases was determined by histological analyses. Tumor infiltrating immune cell profiles were assessed by flow cytometry. Relative distributions of different populations of myeloid cells (C, D, E), and Tregs (F, G) in the tumor microenvironment were determined. IL-17 and Ifng production by the infiltrating T cells in response to PMA and Ionomycin stimulation was measured by intracellular staining (H, I, J). Data displayed in panels A, B, D, E, G, I, and J represent mean±SEM. Significance of differences was determined by One-Way ANOVA with post-hoc Bonferroni test: *P< 0.05; **P< 0.01; ***P< 0.001; ****P< 0.0001.

## DISCUSSION

The tumor microenvironment (TME) is a dynamic assortment of pro-tumor and anti-tumor molecular forces. The success or failure of the establishment and progression of a tumor is dependent on which forces outperform and overpower the others in the TME. Immunosuppressive cell networks and factors play a significant role in the failure of the anti-tumor immune responses and therapies (Ilkovitch and Lopez, 2008). On the other hand, the interferon-gamma (IFNG) cytokine has long been regarded as a marquee orchestrator of the anti-tumor immunosurveillance mechanisms. A recent study suggests that the IFNG transcriptome response in melanoma cells serves to amplify the magnitude of the antitumor T cell response, and the corresponding downstream IFNG signaling factors are the main drivers of the clinical responses to immune checkpoint inhibition (Grasso et al., 2020). Paradoxically, we had previously identified a pro-melanomagenic role of Ifng in the context of UV irradiation in mice (Zaidi et al., 2011). However, it was unclear whether the potential pro-melanomagenic effects of Ifng were due to its systemic functions or via direct stimulation of melanoma cell-mediated events. Our results presented here provide direct in vivo evidence that intracellular Ifng signaling in melanoma cells promotes melanomagenesis and that this promotion is brought about via skewing of the tumor immune microenvironment towards a pro-tumor character.

The specific role of STAT1 in cancer remains ambiguous, as it is known to exhibit both tumor suppressor as well as oncogenic properties in a context-dependent manner (Zhang and Liu, 2017). Several types of cancers, including melanoma, show loss of STAT1, and its tumor suppressor function is mainly attributed to its role in the activation of pro-apoptotic (such as BCL2 and BCL-xL) and cell cycle inhibitor genes (e.g. p27 and p21^WAF1^) as well as negative regulation of angiogenesis (Hsu et al., 2017; Kachroo et al., 2013; Zhang et al., 2014). Several lines of evidence also implicate STAT1 as a key regulator of anti-cancer immunosurveillance by its regulation of the major histocompatibility complex (MHC) Class I and other genes of the antigen presentation machinery (Brucet et al., 2004; Leibowitz et al., 2011; Rodriguez et al., 2007). The most convincing case for the tumor suppressor gene designation for STAT1 is provided by the *Stat1-knockout* mice, which exhibit enhanced susceptibility to spontaneous and carcinogen-induced tumorigenesis (Chan et al., 2012; Lesinski et al., 2003). On the other hand, there is also some evidence for the oncogenic role of STAT1, albeit relatively less convincing (Zhang and Liu, 2017). One possible reason for this dichotomy may rest upon how the analysis of STAT1 expression has been performed in different types of cancers. For example, analysis of the whole tumor tissue fails to distinguish between the STAT1 expression in the tumor cells versus the immune cells. It is plausible to speculate that the expression of STAT1, or lack thereof, may have contrasting effects depending on the cellular context. Our results presented here provide direct experimental evidence that Stat1 plays an oncogenic role in murine melanoma cells.

The *γδ* T cells perform crucial roles in the anti-tumor immune responses, e.g. cytotoxicity, production of IFNG and TNF*α*, and inducing the maturation of dendritic cells (DC). A recent analysis of expression signatures from ~18,000 human tumors with overall survival outcomes across 39 malignancies identified tumor-infiltrating *γδ* T cells as the most significant favorable cancer wide prognostic signature (Gentles et al., 2015). Both positive and negative correlations have been found between clinical responses and tumor-infiltrating *γδ* T cells. The positive correlation between the tumor-infiltrating *γδ* T cells and the clinical survival of the patients was observed in necrotizing choroidal melanomas (Bialasiewicz et al., 1999), ovarian cancer (Raspollini et al., 2005), and melanoma (Cordova et al., 2012). Interestingly, in breast cancer, a potential pro-tumor function was reported (Peng et al., 2007), highlighting that the infiltrating *γδ* T cells were able to inhibit the function of several immune cell populations in vitro and were involved in suppression of anti-tumor responses. Consistent with these observations, the presence of *γδ* T cells was shown to positively correlate with advanced tumor stages and inversely correlated with patient survival. A positive correlation between disease progression and the number of tumor-infiltrating *γδ* T cells was also observed in a cohort of breast cancer patients (Ma et al., 2012). These findings strongly suggest that *γδ* T cells in the tumor microenvironment may play substantially disparate functions; hence positive or negative correlation with prognosis may depend on the specific *γδ* T cell subset present in the TME. In this study, we found that although the frequency of the *γδ* T cells remained approximately the same in the TME, the TME of the tumors made by the Ifng-treated cells harbored mainly CD27-γδ T cells, which have previously been shown to produce IL-17 rather than Ifng (Ribot et al., 2009). We found that the Ifng-treated melanoma cells induced a tumor microenvironment that was enriched in pro-tumorigenic IL-17, TGFβ-producing γδ T cells, and accumulation of immunosuppressive polymorphonuclear leukocytes (PMN). We also demonstrated that the genetic ablation of γδ T cells not only reduced the tumor burden and metastasis but also substantially reduced the accumulation of PMN, thus providing direct evidence of the involvement of pro-tumorigenic γδ T cells in PMN accumulation and melanoma progression and metastasis. Most interestingly, ablation of the γδ T cells could also restore anti-tumor immune response in the Ifng-treated melanoma microenvironment as evidenced by an increased frequency of Ifng-producing CD4+ and CD8+ T cells and striking down-modulation of CD25 and Foxp3 expression in the T_regs_ akin to ‘non-T_reg_’ (Togashi and Nishikawa, 2017).

Our results suggest that IFNG is the driver of novel cellular/molecular inflammatory mechanisms that may underlie the outgrowth of melanoma. Melanocytes are built for enhanced survival, to withstand both UV exposure ensuring the continued synthesis of melanin, and the chemical stresses associated with the presence of melanin itself. The microenvironmental elements in the aftermath of UVR insult may play an integral role in further protecting melanocytes from eradication by the UVR-induced inflammatory microenvironmental response. Melanoma can take advantage of this built-in circuitry to develop into one of the most evasive cancers. The fact that this circuitry converges on IFNG signaling epitomizes the importance of this pathway to melanocytic survival mechanisms. While it adds further complexity to the already intricate melanocytic microenvironmental interactions, it also offers an opportunity to understand the process of melanoma progression from a new perspective. At the same time, it promises to identify potentially novel targets for both the prevention and therapy of melanoma. For example, topical inhibition of IFNG signaling in the immediate aftermath of sunburn may be explored as a means to prevent melanocyte activation and an enhanced eradication of UVR-damaged melanocytes, reducing susceptibility to malignant transformation. Moreover, inhibition of IFNG signaling may enhance the efficacy of immunotherapy against melanoma as well as other cancers, e.g. antibody-mediated blockade of CTLA4 and PD-1/PD-L1 (Callahan et al., 2013; Drake et al., 2014).

## METHODS

### Cell lines

Five mouse melanoma cell lines were used in this study. B16 cell line was obtained from Dr. Glenn Merlino (NCI/NIH). B16N is a novel metastatic clone of B16, established at NCI. B16 and B16N both are syngeneic to the C57BL/6 strain background. The B2905 (C57BL/6) and F5061 (FVB/N) cell lines were derived from spontaneous tumors induced by UV irradiation of the hepatocyte growth factor/scatter factor (HGF/SF) transgenic mice (Noonan et al., 2001; Noonan et al., 2012). YUMM1.1 cell line was isolated from a Braf^V600E^; Pten^−/−^; Cdkn2a^−/−^ mouse melanoma was obtained from Dr. Marcus Bosenberg (Meeth et al., 2016). Human melanoma cell line A2058 was purchased from ATCC (CRL-11147). All cell culture media and supplements were purchase from Life Technology. All tumor cell lines were cultured at 37°C in DMEM supplemented with 10% FBS, l-alanyl-l-Glutamine (2 mmol/L), and Gentamycin (50 μg/mL) at 5% CO_2_. DMEM, FBS, and l-alanyl-l-Glutamine were purchased from Corning, Cellgro.

### Generation of CRISPR/Cas9-mediated knockout cell lines

We designed two guide RNAs targeting different exons of Stat1 and Stat3 loci by online CRISPR Design Tool. The Cas9 expression construct pSpCas9(BB)-2A-GFP was purchased from Addgene (Plasmid ID 44758). Stat1 (NM_001205313.1) and Stat3 (NM_213659) were used to search gRNA using the online CRISPR Design Tool (http://tools.genome-engineering.org) (Ran et al., 2013).Two different gRNA targeting different exons were used for both Stat1 and Stat3. The sequence of gRNA_Stat1_#1: GGAAACTGTCATCGTACAGC. The sequence of gRNA_Stat1_#2: GGTCGCAAACGAGACATCAT. The sequence of gRNA_Stat3_#1: GCAGCTGGACACACGCTACC. The sequence of gRNA_Stat3_#2: TTCTTCACTAAGCCGCCAAT. Plasmid construction and molecular cloning were done by following the previously published protocol (Ran et al., 2013). B16N cells were transfected with each construct using Lipofectamine 3000 (Invitrogen) following the manufacturer’s protocol. Single GFP+ cells were sorted into each well of multiple 96-well plates by BD Influx Cell Sorter after 48 h post-transfection. Selected clones were screened for expression of either Stat1 or Stat3 by quantitative real-time PCR, western blotting, and Surveyor mutation detection assay.

### Mice

The C57BL/6, FVB/N, athymic Nude, *Ifng-knockout* (*Ifng*^*tm1Ts*^), and *B6-TCRdelta-knockout (*<u>*Tcrd*^*tm1Mom*^</u> *allele*), *B6-TCRbeta-knockout (Tcra*^*tm1Mom*^ *allele*) *mice* were purchased from The Jackson Laboratory. The NOD.Cg-*Prkdc*^*scid*^ *Il2rg*^*tm1Sug*^/JicTac (CIEA NOG) immunodeficient mice were purchased from Taconic Biosciences. Both female and male mice were used at 6-8 weeks of age, with individual experiments using mice of a single sex.

### Antibodies

All fluorescently labeled antibodies used were obtained from commercial sources (eBioscience or Biolegend), including anti-Thy1, TCRβ, γδTCR, CD4, CD8α, CD11b, CD19, Ly6g (1A1), CD45.2, Foxp3, CD25, GITR, PD1, CD27. CCR6 antibody was purchased from R&D.

### Interferon-gamma treatment of cells

Recombinant mouse Ifng and human IFNG proteins (with carrier) were purchased from Cell Signaling Technology (catalog #39127 and 8901). They were reconstituted with sterile water at a concentration of 0.1 mg/ml, then diluted into a final concentration of 10 ng/ml with fresh DMEM with 10% FBS. According to safety data sheets provided by the manufacturer, the bioactivity of h-IFNG was determined in a virus protection assay. The ED50 of each lot was between 0.3 and 1.2 ng/mL. The conversion of 10 ng/mL to biological activity was 8.33 to 33.33 U/mL. The cells were treated with 10 ng/ml Ifng/IFNG in culture at 50% confluency for 48 h followed by 3x washes with PBS just before inoculation in mice and other assays.

### Tumor cell inoculation and tumor analysis

All animal studies were approved by the Institutional Animal Care and Use Committee at Temple University. Melanoma cells with or without GFP-luciferase reporter were cultured and counted using an automated cell counter (BioRad). Single cell suspension in 100 μL of 1×PBS were transplanted subcutaneously to mice or introduced by tail vein injection with a 1-ml syringe with 30 (½)-gauge needle. *For subcutaneous injection:* For B16, B16N, and B2905 cells, 2.5×10^5^ cells were injected; for F5061 and YUMM1.1 cell, 1×10^6^ cells were injected in flanks. Tumor latency was defined as the period between injection of tumorigenic cells into mice and the appearance of tumors of ≥1 mm in diameter. The endpoint was a tumor diameter of 1 cm. Tumor growth was measured by caliper, and tumor volumes were calculated using the formula ½*(L × W× H)*. *For tail vein injection:* For B16, B16N, and B2905 cells, 1.25×10^5^ cells were inoculated through the tail vein. 5×10^5^ cells were injected for F5061 and YUMM1.1 cells. All cell lines were injected in mice on the C57BL/6 strain background, except for the F5061, which were injected in FVB/N mice. Bioluminescent imaging (BLI) was performed in a Xenogen IVIS imaging system (Perkin Elmer) after intraperitoneal injection of luciferin (100 μl of 15 mg/ml solution per 10 g). The endpoint was the day of euthanasia as determined by >10% body weight loss, hind limb paralysis or fracture, immobility, or a total photon flux > 1 × 10^8^, a value that initial results indicated reliably predicts death in less than one week in this model. Lungs and other distant organs (such as liver, spleen, kidney, right femur for bone marrow collection, brain, and any observed potential metastatic tissues) were removed from mice at day 21 post-injection. Some samples were collected for flow cytometry, while others were perfused and fixed in 4% paraformaldehyde (Fisher Scientific) overnight at 4°C, rinsed, and transferred to 30% ethanol, and stored at 4°C until further analysis. Lung surface gross tumor nodules were counted under a dissecting microscope. Images of whole mouse lungs were captured using a Nikon SLR camera with AF 60mm 1:2.8 D lens (magnification 1×).

### Preparation of single cell suspensions for flow cytometry

Tumors were micro-dissected from mouse tissues at the end of each experiment, dissociated through a 50-μm filter, and washed with PBS. In some cases, immune cells were further enriched by layering cell suspension γδ on the Ficoll–Hypaque, followed by centrifugation for 15–30 min at 400 g. The buffy layer was isolated and washed twice with RPMI before staining and FACS analysis. To measure cytokine production, cells were stimulated with PMA/ionomycin in presence of Brefeldin A for 6 h before intracellular staining for FACS. analyses.

### Intracellular Staining

Cells were stained for surface markers, then fixed in 100 μl of Cytofix/Cytoperm solution for 30min at 4°C, washed 2X two times in Perm/Wash solution, pelleted by centrifugation and resuspended in 100 μl of Perm/Wash solution with or without (FMO control) fluorochrome-conjugated antibody at room temperature, using the BD Permeabilization Solution obtained from Transcription Factor Phospho Buffer Set (Cat. No. 565575), according to manufacturer’s instructions.

### Fluorescence-activated cell sorting

B16N melanoma cells were transfected with a Green Fluorescent Protein (GFP) and Firefly Luciferase lentivirus after single-cell growth selection process. GFP-positive cells were sorted using the BD FACSAria IIu or BD FACS Vantage (BD Biosciences) systems. FACS DIVA software was used during cell sorting and the FlowJo software for analysis. Cells were initially identified on forward scatter (FSC) vs. side scatter (SSC). Single cells were identified using FSC and SSC pause width. Cell doublets were excluded from the analysis. Untransfected cells were used as negative controls. Cells were sorted based on GFP expression and SSC-A. GFP-positive cells were identified using appropriate gates based on negative controls. Due to low sample cell number, reanalysis of sorted cells was not usually done, but representative post-sort analyses confirmed that presort purities of 0.74–0.75% were enriched to 98–99.5%.

### Western blotting

Human melanocytes and melanoma cells were lysed in Pierce RIPA buffer (Thermo Scientific) containing 1× Halt protease inhibitor cocktail (100×; Thermo Scientific) and 1x Halt phosphatase inhibitor cocktail (100×; Thermo Scientific), and the protein concentration was measured with the Bio-Rad Protein Assay following the manufacturer’s protocol. The same amounts of protein extracts were subjected to polyacrylamide gel electrophoresis using the 4% to 20% Mini-Protean TGX gel system (Bio-Rad), transferred to PVDF (0.45 μm pore size; Millipore) membranes, and immunoblotted using antibodies that specifically recognize STAT1 (1:1,000; Cell Signaling Technology), pSTAT1 (Y701, 58D6, 1:1,000; Cell Signaling Technology), pSTAT1 (Y727, D3B7, 1:2,000; Cell Signaling Technology), STAT3 (124H6, 1:1,000; Cell Signaling Technology), pSTAT3 (Y705, D3A7, 1:2,000; Cell Signaling Technology), GAPDH-HRP (D16H11, 1:1,000; Cell Signaling Technology), IRF1 (D5E4, 1:1,000; Cell Signaling Technology). The secondary antibodies used for detection were horseradish peroxidase (HRP)–conjugated goat anti-mouse and goat anti-rabbit IgG (1:5,000; Thermo Scientific). The blots were incubated with Luminata Western HRP substrate (Millipore) for 5 minutes. Band intensities of Tiff images were quantified by using Image J software.

### Quantification and statistical analyses

All sample sizes were determined based on preliminary studies and prior knowledge of expected variability within assays. For animal studies, age-matched (6–8 weeks) mice were randomly assigned to control and experimental groups. Quantification of the lung nodule counts were performed blindly by the pathologist. Where blinding was not used, data were analyzed using automated image analysis software when possible. All statistical tests used were deemed appropriate and met the assumptions required; when parametric tests were used, normal distribution was assumed. Where necessary unequal variance was corrected for, or if no correction was used, variation was assumed equal based on prior knowledge of the experimental assay. All experiments were performed in triplicate, data are presented as mean ± SEM, and graphs were prepared with GraphPad Prism 7. To analyze the statistical difference between two groups, a two-tailed unpaired Student’s t-test was used. Comparisons involving multiple groups were assessed by one-way ANOVA with post-hoc Tukey analysis. P value < 0.05 was considered as statistically significant.

### Cell proliferation assay

A total of 3,000 cells in 200 μl of DMEM plus 10% FBS was plated in 6 wells per cell type per condition in a 96 well plate. To measure cells input on day 0, an additional set of 6 wells per cell type was seeded and assayed 2 hours later. Plates were developed by discarding 100 ul from each well and adding 100 ul of solution containing DMEM with 10% FBS and WST-1 reagent (Roche Diagnostics) diluted 1:10. Plates were incubated for 1 h at 37°C and absorbance at 450 nm was measured using a plate reader.

### *In vitro* Tumor Growth Assays

Neutralized rat tail collagen solution was prepared at 0.8 mg/ml in DMEM by adding appropriate amounts of 10× DMEM concentrate and 1 N NaOH. Next, melanoma cells were resuspended at 2.86 × 10^4^ cells/ml in the collagen solution, and 350 μl of cell suspension was plated per well in 24 well plates (for a final cell number of 10,000 cells per well). After 20 min at 37 °C, wells containing cells, suspended in polymerized collagen, were overlaid with 3,000 tumor cells per well in 500 μl of DMEM with 10% FBS. Tumor cell growth was photographed, and images were analyzed by Image J software.

### Soft-Agar Colony Formation Assay

A soft-agar colony formation assay was done using 6-well plates. Each well contained 2 mL of 0.8% agarose base layer in complete medium (DMEM with 10% fetal bovine serum and 1% antibiotics) as the bottom layer and 1 mL of 0.4% agarose in complete medium and 3,000 cells (untreated and Ifng treated cells) as the top layer. Cultures were maintained under standard conditions for 14-21 days. The colonies were stained with cell stain solution (Chemicon) overnight at 37°C and counted the following morning. The number of colonies was determined with a microscope at ×100 magnification; a group of >20 cells were counted as a colony. For colony quantification 1.4 mL cell quantification solution (Chemicon) was added to each well and incubated for 4 h at 37°C. Absorbance was measured at 490 nm.

## ACKNOWLEDGEMENTS

This work was funded by USA National Institutes of Health, R01CA193711 (MRZ); R01CA236391 (DJK), R01AI068907 (DJK), P30CA006927 (FCCC Comprehensive Cancer Center Core Grant).

## AUTHOR CONTRIBUTIONS

BZ and JB: Conceptualization, methodology, investigation, data acquisition, and manuscript writing. HRK, XM, and SPA: methodology and data acquisition. KQC: Histopathological analyses and quantification. DJK: Investigation, acquisition of the grant, data interpretation, review and editing of manuscript. MRZ: Conceptualization, acquisition of grant, data interpretation, project supervision, and manuscript writing.

## CONFLICTS OF INTEREST

Authors have no conflicts of interest.

---

**Figure S1.**
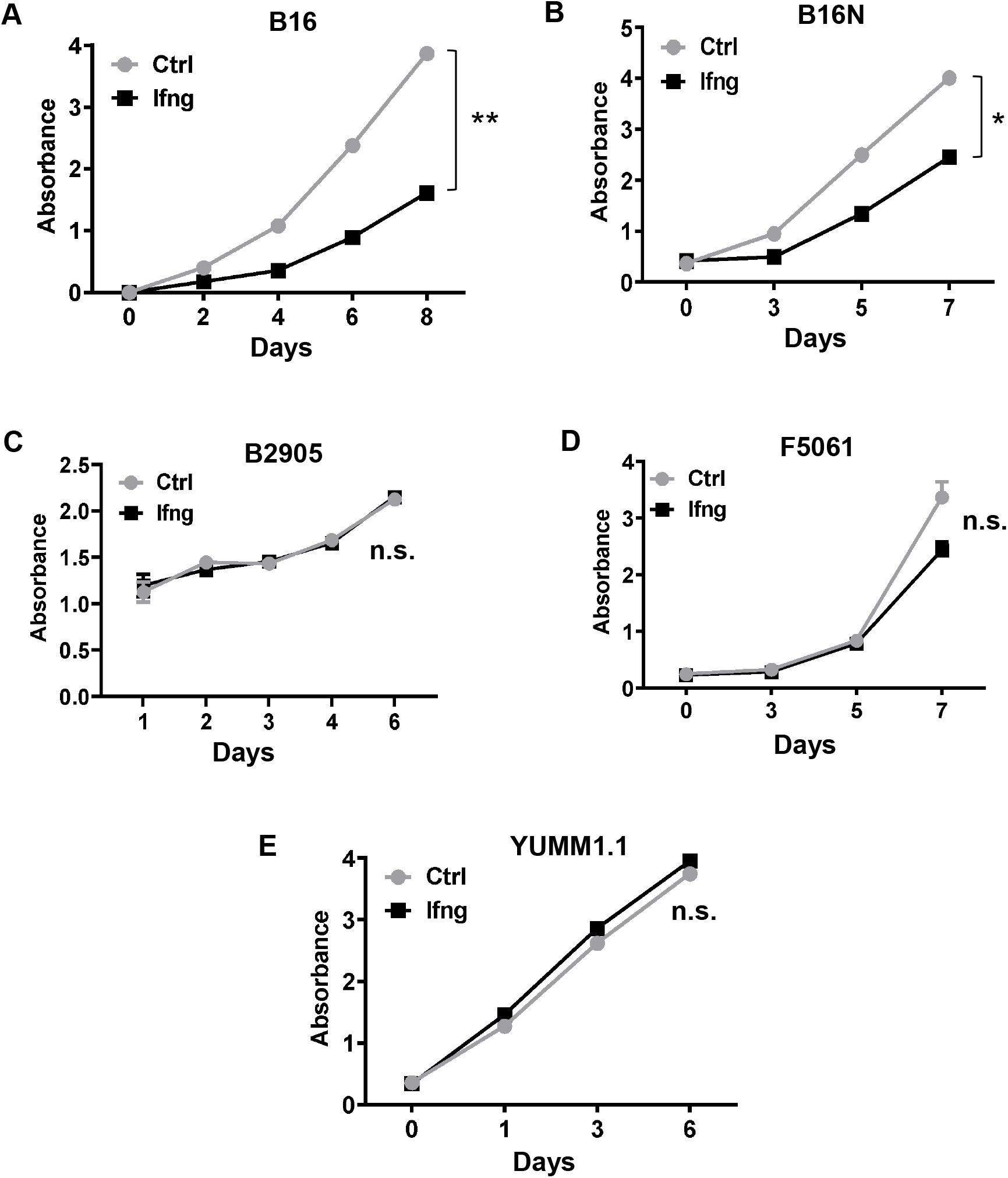
Ifng treatment showed differential effects on cell proliferation in five indicated mouse melanoma cell lines. The proliferation of (A) B16 and (B) B16N cell lines was significantly inhibited. In contrast, (C) B2905, (D) F5061, and (E) YUMM1.1 cell lines were not affected. For proliferation, WST-1 assay was performed with mock- or Ifng treatment with 10 ng/ml conc. Melanoma cells in medium with or without Ifng were seeded in triplicates and absorbance was measured at indicated time points. All data are plotted as mean±SEM. *P<0.05; **P<0.01; n.s. not significant; Student’s t test.

**Figure S2.**
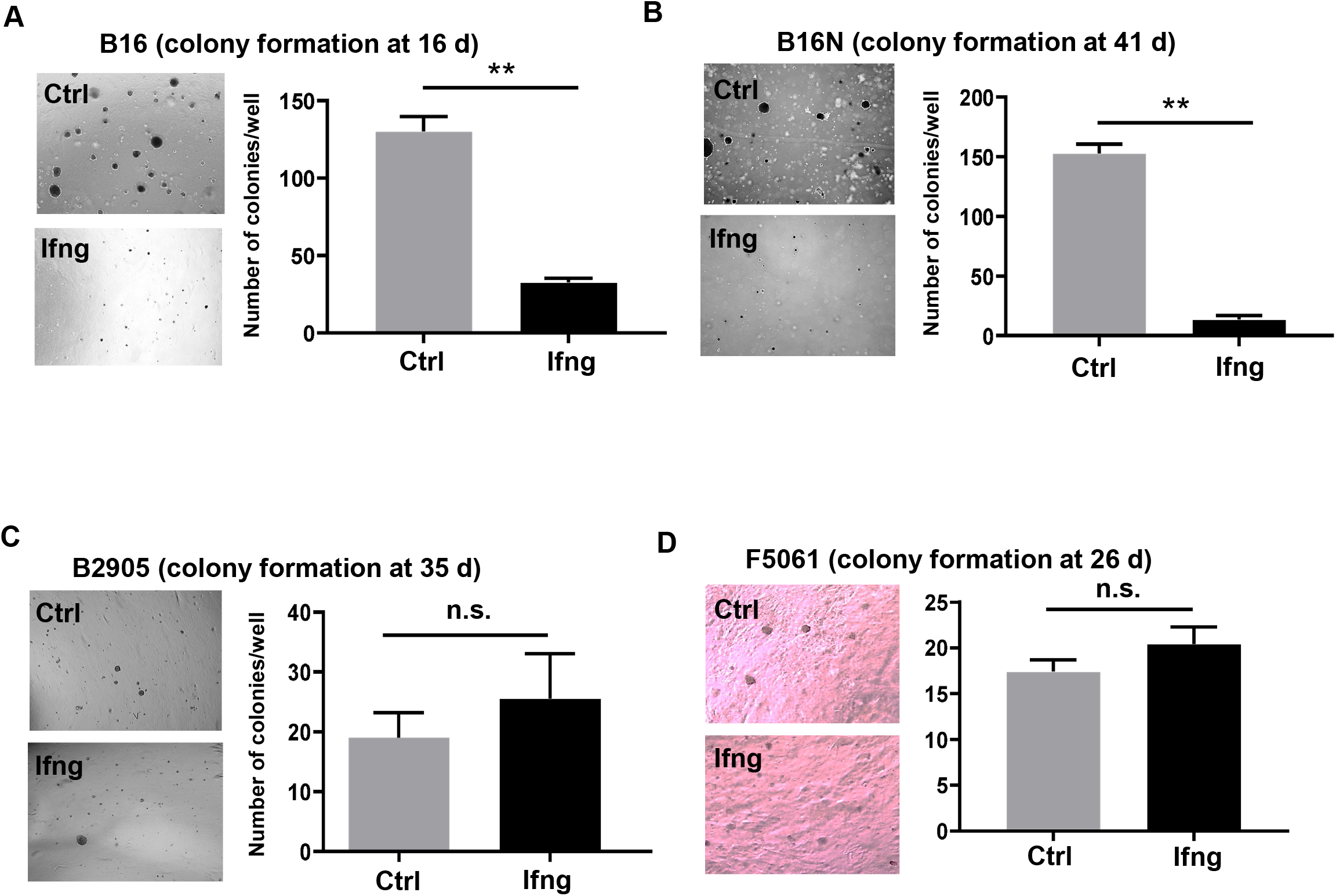
Ifng treatment showed differential effects on soft agar colony formation in four indicated melanoma cell lines. (A) B16 and (B) B16N cell lines were significantly inhibited in colony formation by Ifng. In contrast, (C) B2905 and (D) F5061 cell lines were not affected. The number of colonies per well were counted after indicated number of days of incubation. Representative microscopic images are shown. All experiments were performed in triplicates and plotted as mean±SEM. **P<0.01; n.s. not significant; Student’s t test.

**Figure S3.**
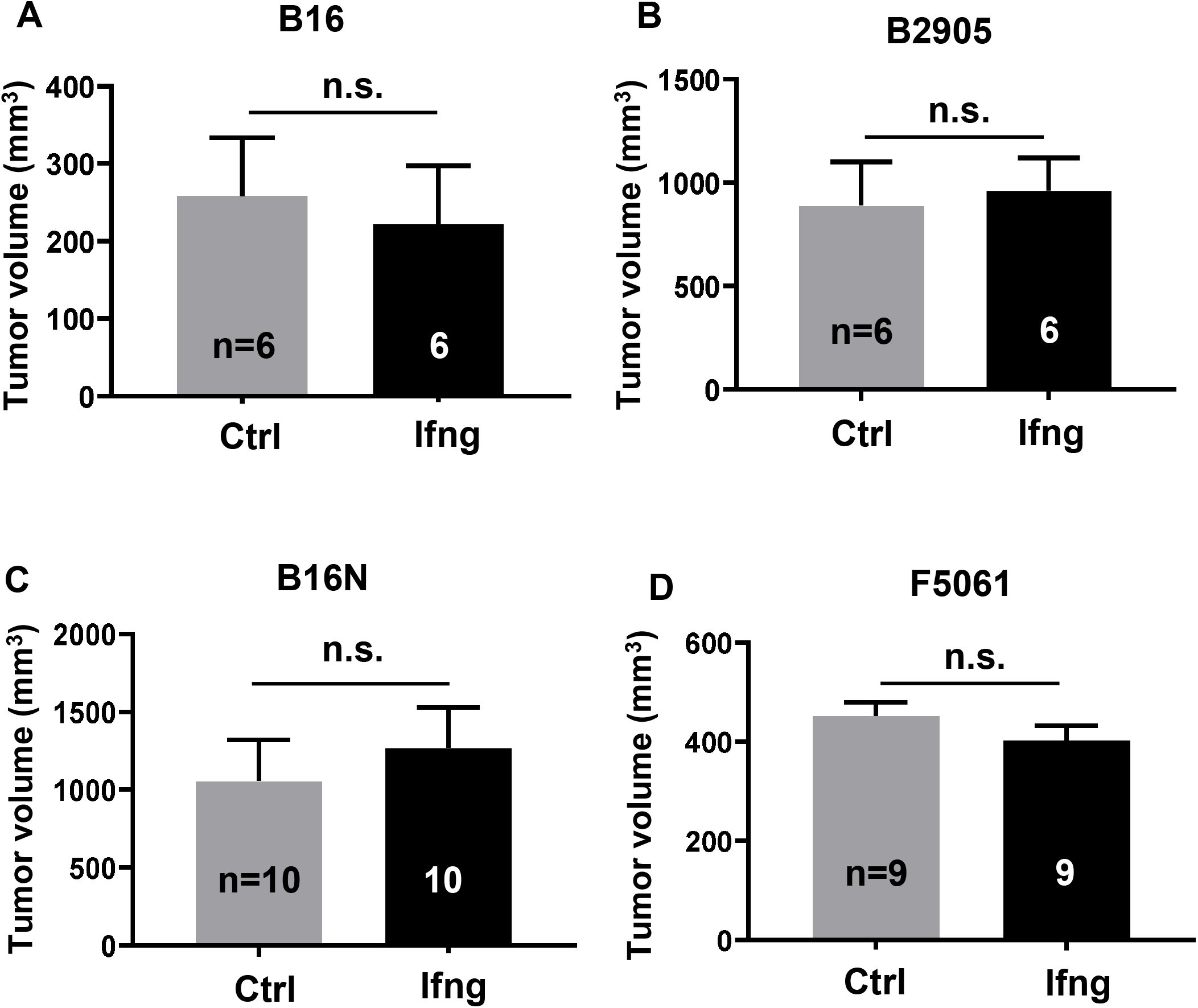
Ifng treatment did not affect subcutaneous growth of melanoma cells. The mouse melanoma cells were mock- or Ifng-treated (10 ng/ml for 48 h) and subcutaneous injections were performed at back flank of 8-week-old syngeneic mice (C57BL/6 for B16, B16N, and B2905; FVB/N for F5061). Once tumors became palpable, tumor volumes were measured up to 3x a week with digital calipers. There was no significant difference in mean subcutaneous tumor volume between control and Ifng-treated (A) B16, (B) B2905, (C) B16N, and (D) F5061 cells. Data are presented as mean±SEM; n.s. not significant, Student’s t test.

**Figure S4.**
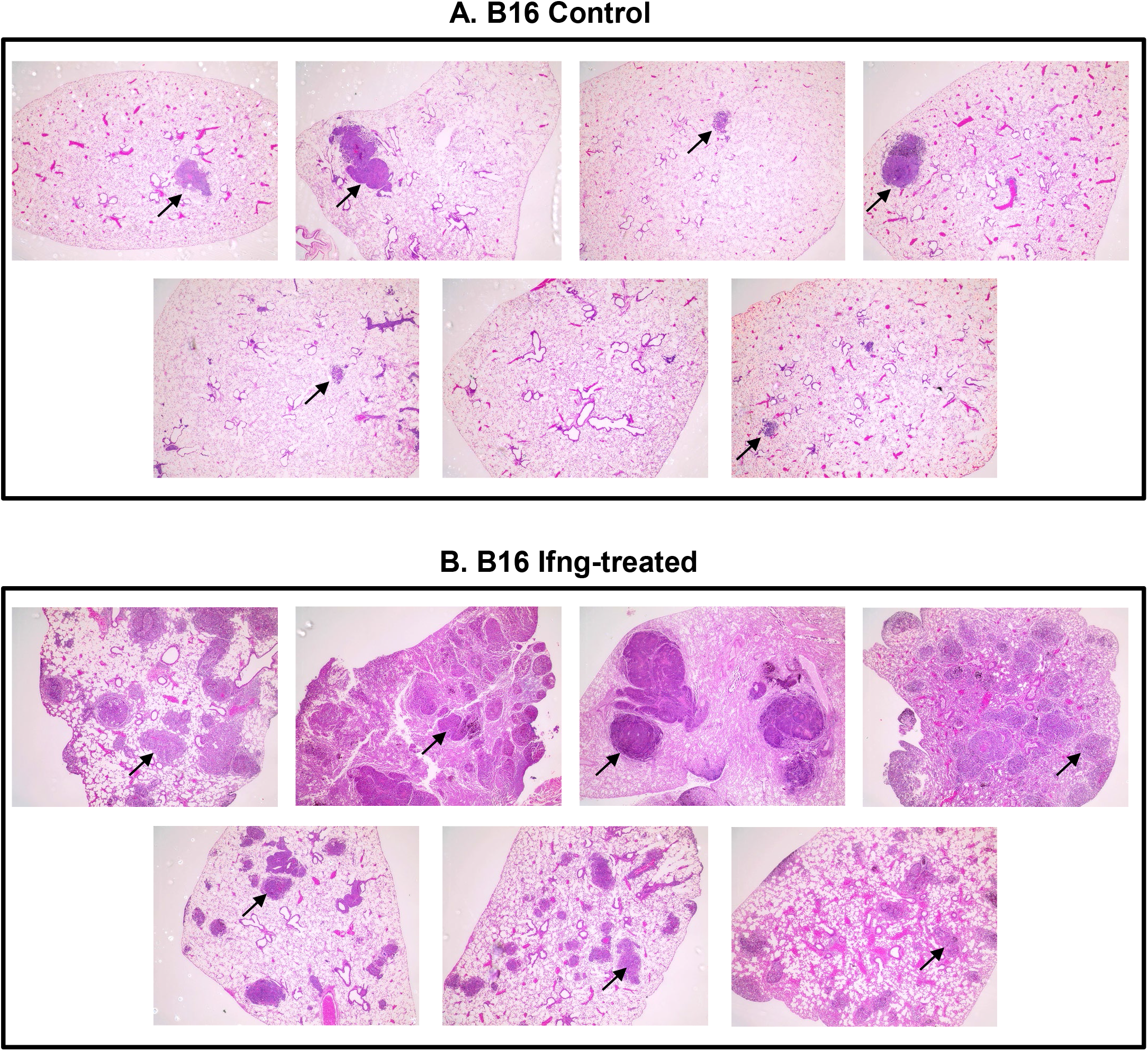
H&E stained FFPE sections for lung tissues from mice injected via tail-vein with (A) mock-treated control or (B) Ifng-treated B16 cells. Cells were treated with 10 ng/ml mouse recombinant Ifng in culture for 48 h, followed by 3x washes with PBS and injected into syngeneic C57BL/6 mice via tail vein (n=7 each). Tumors are seen as darker purple lesions. Only a few random lesions are marked by arrows, especially in (B).

**Figure S5.**
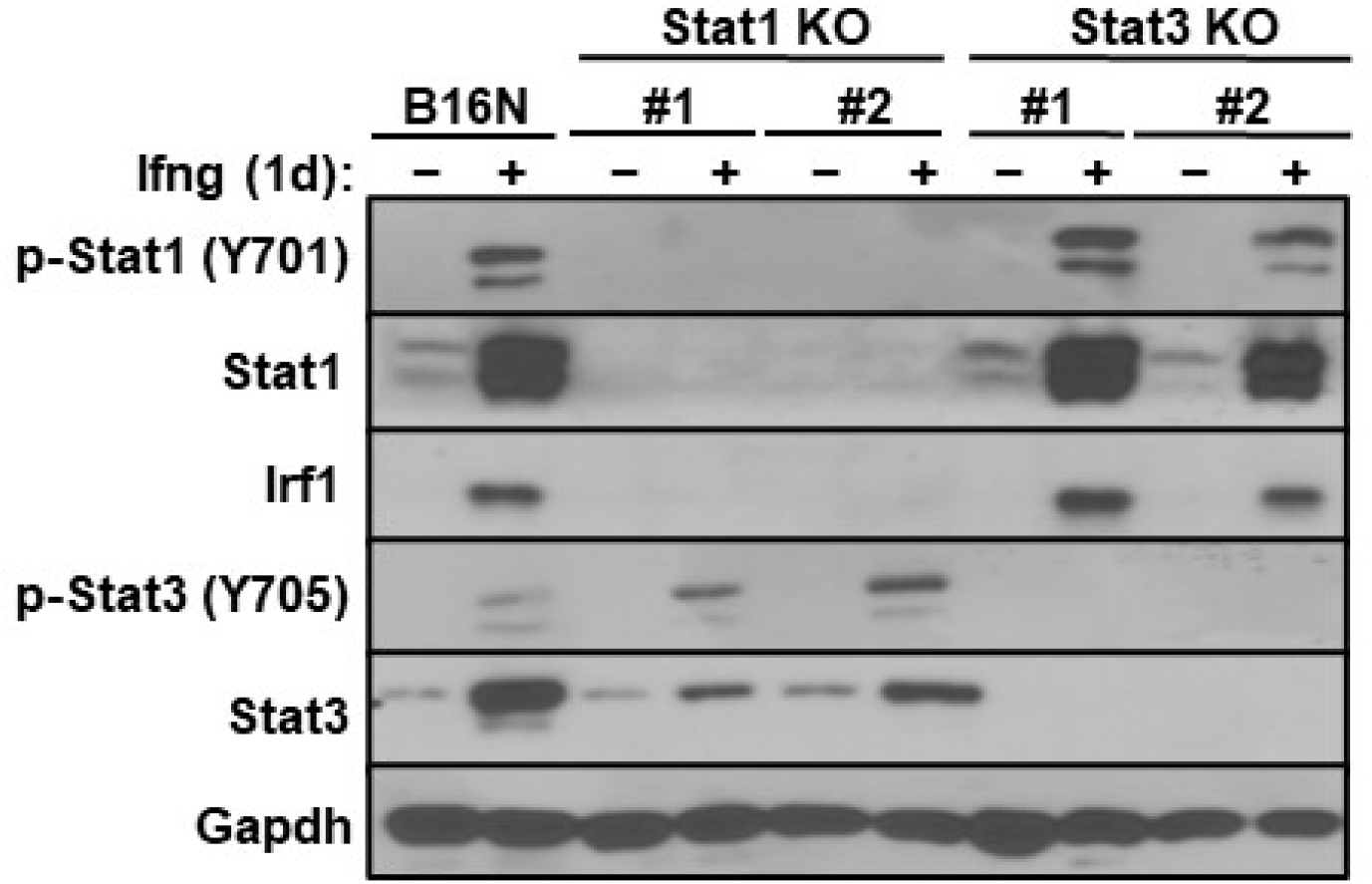
Western blot of pStat1 (Y701), Stat1, Irf1, pStat3 (Y705), Stat3, and Gapdh expressions in parental B16N cells, Stat1-KO and Stat3-KO clones (2 clones each, #1 and #2) in response to 10 ng/ml Ifng treatment for 24 h.

**Figure S6.**
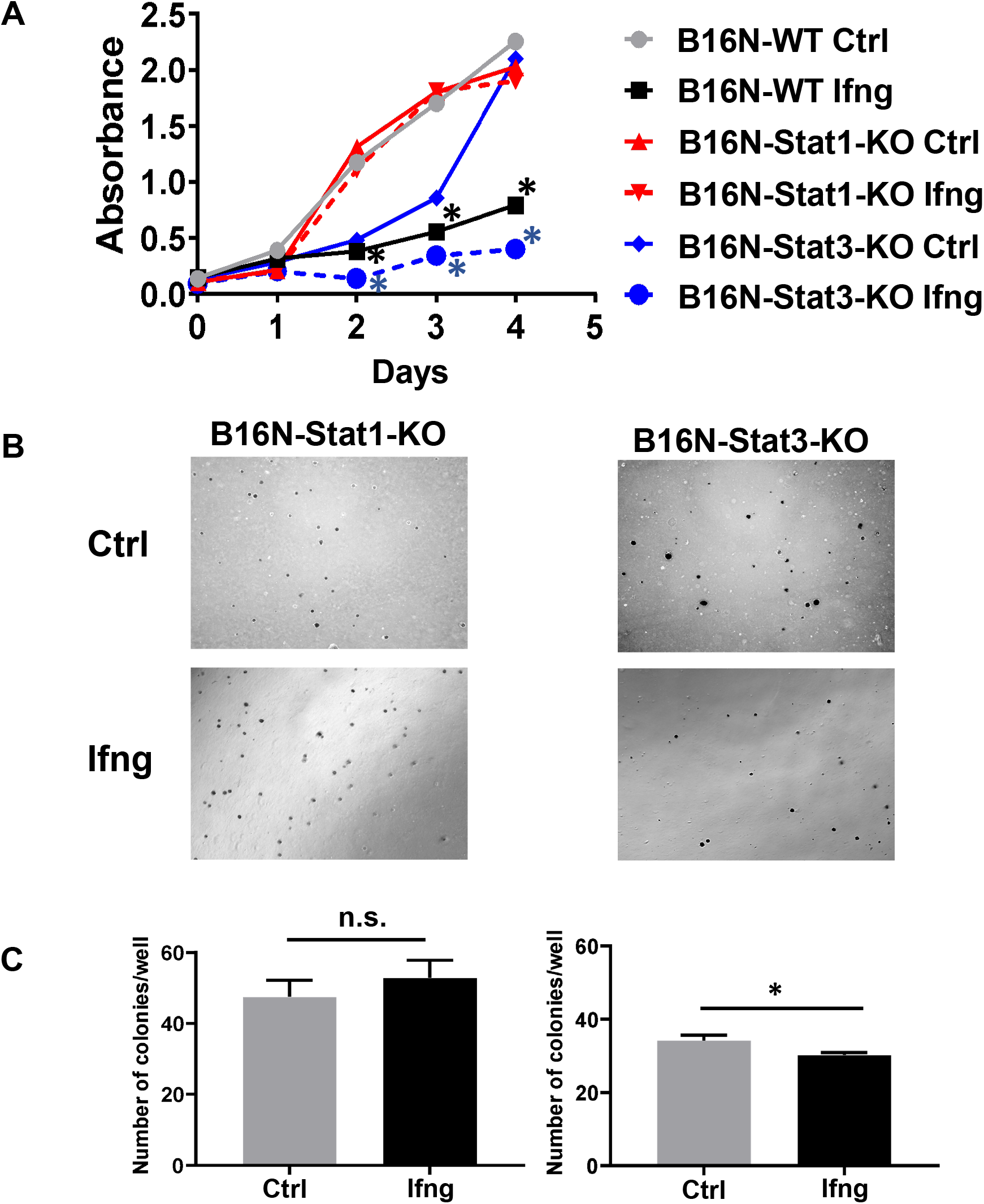
The effect of Ifng on B16N cell proliferation and colony formation is dependent on Stat1 but not Stat3. **(A)** Ifng treatment inhibited the growth of Stat3 knockout (KO) and parental B16N cells, but not Stat1-KO cells. Cell proliferation assay (WST-1) showed an Ifng-induced reduction of cell growth on WT and Stat3-KO cells (both P<0.05 on day 2 onward), but not Stat1-KO cells. Six replicates per sample were measured and plotted as mean±SEM. Plates were read at 450nm for absorbance after 4 days culture. **(B)** Ifng had no effect on colony formation on Stat1-KO but had reduced colony formation on Stat3-KO cell lines. The number of colonies per well for Stat1-KO and Stat3-KO cells in soft agar assay with representative microscopic images are shown. Images were taken at day 27 (Magnification 4×). **(C)** Quantified colonies data are plotted as mean±SEM of three independent experiments. *P<0.05; n.s. not significant, Student’s t test.

**Figure S7.**
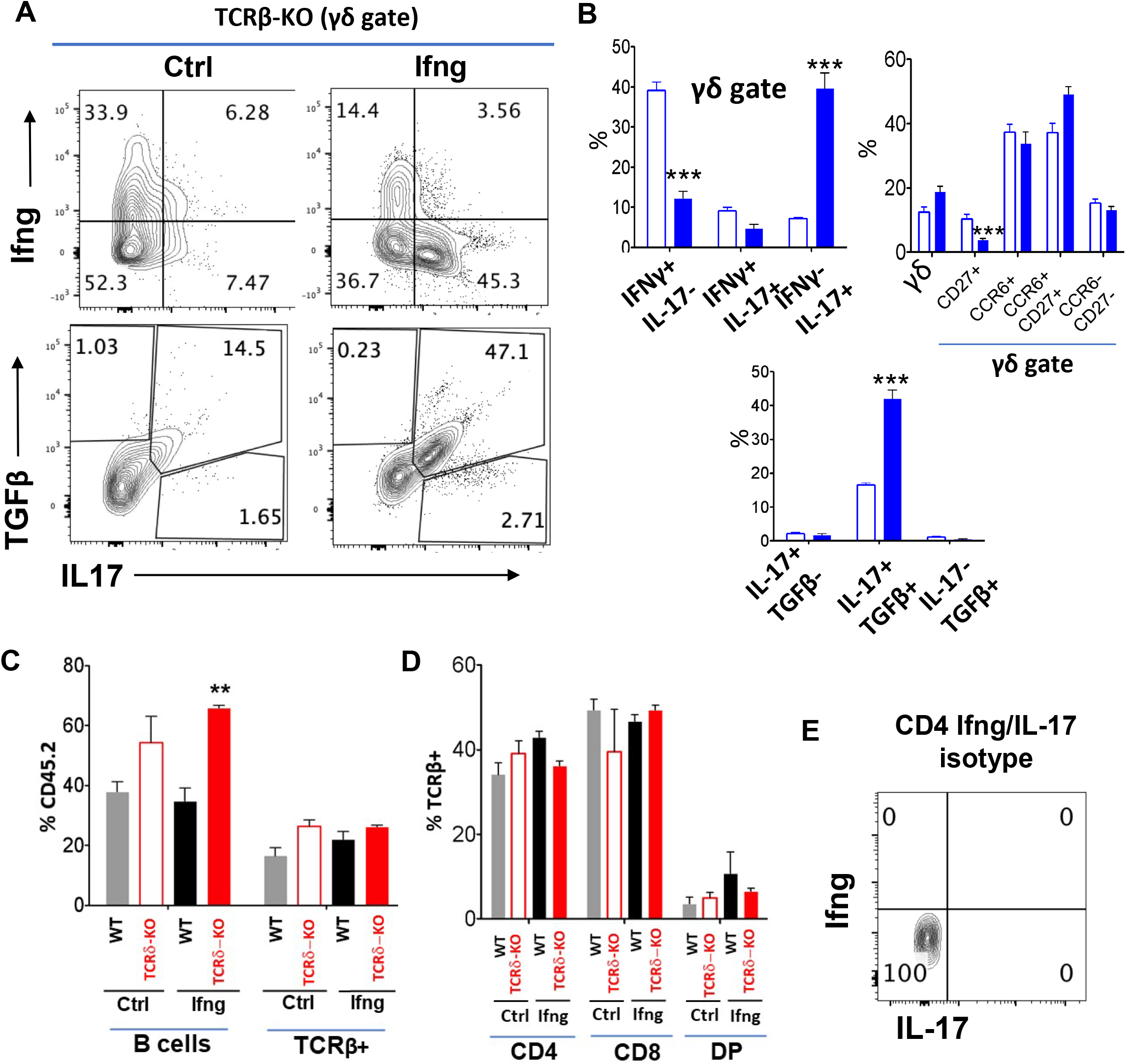
Requirement for gamma-delta T cells in Ifng-enhanced metastasis. Ifng treated and untreated B2905 melanoma cells were inoculated into syngeneic WT, TCRβ−/− and TCRγ−/− mice by tail vein injection. Mice were euthanized on day 23. **(A)** Tumor infiltrating immune cell profiles were assessed by flow cytometry. **(B-D)** Relative distributions of different populations of myeloid cells and Tregs in tumor microenvironment were determined. **(E)** IL-17 and Ifng production by infiltrating T cells in response to PMA and Ionomycin stimulation was measured by intracellular staining. Data displayed represent mean ± SEM. Significance of differences was determined by one-way ANOVA with post-hoc Bonferroni test, as indicated by asterisks: * p < 0.05; ** p < 0.01; *** p < 0.001; **** p < 0.0001.

## Notes

### Competing Interest Statement

The authors have declared no competing interest.

